# Immune Cell Influence on Diffuse Large B-Cell Lymphoma: A Mendelian Randomization Analysis

**DOI:** 10.1101/2024.09.05.611435

**Authors:** Honghua He, Jihong Zhong, Qinghua Li, Chen Deng, Xin Yuan, Kaixiang Zhang, Lirong Nie, Nali Cai

## Abstract

**Objective:** To elucidate the causal relationship between immune cells and diffuse large B-cell lymphoma (DLBCL), we conducted a Mendelian randomization analysis.

**Methods:** Mendelian randomization (MR) leverages genetic variants as instruments to infer causal effects from observational data. Here, we performed a two-sample MR analysis to assess the causal impact of 731 immune cell types on DLBCL. We employed various MR techniques, including the weighted median estimator (WME) and inverse variance weighting (IVW), and conducted sensitivity analyses to ensure result robustness. Additionally, reverse MR analysis was performed to explore the potential causal relationship between DLBCL and immune cells.

**Results:** We identified seventeen immune features with causal links to DLBCL, categorized across various cellular groups: four in B cells, two in T cell maturation stages, six in Tregs, four in the TBNK group, and one in dendritic cells (DCs). Sensitivity analyses confirmed the absence of heterogeneity, horizontal pleiotropy, and bias in our findings. Reverse causal analysis revealed a causal association between DLBCL and one of the seventeen immune cell types identified.

**Conclusions:** Our MR analysis of seventeen immune cell types uncovers the complex interactions between the immune system and DLBCL, providing crucial insights into the tumor microenvironment and potential avenues for targeted immunotherapy.

## 1 Introduction

Diffuse large B-cell lymphoma (DLBCL) is an aggressive tumor derived from mature B lymphocytes[1]. As the most prevalent type of non-Hodgkin lymphoma (NHL), it is distinguished by its diffuse growth pattern and widespread destruction of nodal architecture or extranodal infiltration by large B lymphoid cells[2]. DLBCL represents 30-58% of all NHL cases worldwide, with an annual incidence rate of 3.8 per 100,000 in Europe[3]. Genetic alterations and environmental signals disrupt the regulatory landscape, resulting in the loss of normal physiological states and the emergence of stable disease states such as DLBCL[4]. DLBCL is a genetically diverse disorder characterized by numerous low-frequency mutations, somatic copy number alterations (SCNAs), and structural variants (SVs)[5]. The microenvironment of B-cell lymphoma (LME) is complex and includes blood vessels, extracellular matrix, stromal cells, and a diverse array of immune cells such as T cells, macrophages, and natural killer (NK) cells[6]. DLBCL cells frequently utilize mechanisms to evade immune surveillance, including the upregulation of immune checkpoint molecules like PD-L1[7].PD-L1 is expressed on immunosuppressive cells within the tumor microenvironment and in a subset of DLBCL case[8-11]. The engagement of PD-1 with its ligands appears to induce cell death and suppress signaling in lymphocytes and monocytes[12].

Previous studies from high-income countries have demonstrated that peripheral blood monocyte count[13-15], lymphocyte-to-monocyte ratios (L:M)[16,17], and lymphocyte-to-neutrophil ratios (N:L)[18,19] are prognostic for DLBCL. Additionally, the numbers of regulatory T cells (Tregs) and HLA-DR low-expressing monocytes[20], also known as immunosuppressive monocytes, have been linked to prognosis. However, reports on peripheral blood Treg numbers in DLBCL have been conflicting: a study from Taiwan found significantly higher Treg counts in DLBCL patients compared to healthy controls [21], whereas a study from Poland observed lower Treg levels [22]. These discrepancies may stem from confounding factors, reverse causation, and sample size limitations, which can introduce bias into observational studies[23]. DLBCL treatment has markedly improved over the past few decades, particularly with the advent of immunochemotherapy regimens such as R-CHOP (rituximab, cyclophosphamide, doxorubicin, vincristine, and prednisone)[24]. Around 60% of patients attain long-term remission with first-line treatment[25]. Nevertheless, about one-third of DLBCL patients experience refractory disease or relapse, presenting significant therapeutic challenges and underscoring the need for novel targeted and immunotherapeutic strategies[26-28].

Mendelian randomization (MR) uses genetic variants as instrumental variables to infer causal relationships between exposures and outcomes, thereby minimizing the effects of confounding and reverse causation common in traditional observational studies[29]. Utilizing genetic variants linked to immune cells, MR provides a robust method to determine whether these cells causally contribute to DLBCL development. This approach not only refines causal inference but also aids in identifying potential therapeutic targets and enhancing risk prediction[30]. Applying MR analysis to DLBCL research can deepen our understanding of the disease’s pathogenesis and guide the development of more personalized and effective clinical strategies.

This study employed Mendelian randomization to perform a comprehensive two-sample bidirectional analysis of 731 immune cell phenotypes and DLBCL, aiming to reveal potential causal relationships between these immune cell phenotypes and the disease(Figure 1).

**Figure 1:**
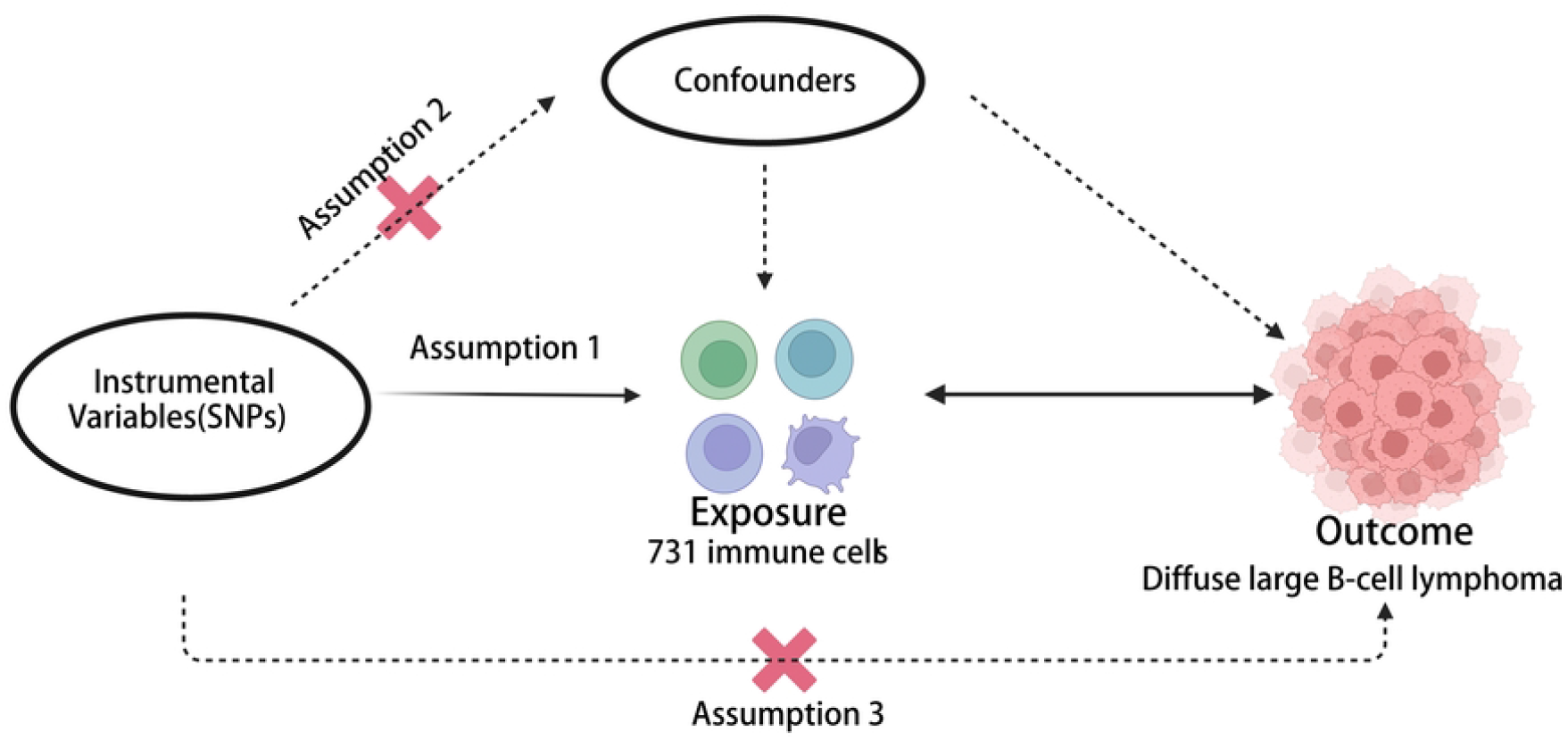
Overall design of the study.

## 2 Materials and Methods

### 2.1. Study design

In this study, we performed an extensive evaluation of 731 immune cell types as potential exposure factors, using single nucleotide polymorphisms (SNPs) associated with these cells as instrumental variables. DLBCL was chosen as the primary outcome variable. We applied a robust two-sample Mendelian randomization approach to analyze the data, carefully assessing for heterogeneity and pleiotropy, and conducting sensitivity analyses to ensure result reliability. The significant immunophenotypes identified in the forward MR analysis will be used as outcome variables in subsequent analyses to explore the causal relationship between DLBCL and immune cells.

### 2.2. GWAS data sources

We obtained data on 731 immune cell types from the GWAS Catalogue (https://www.ebi.ac.uk/gwas/home), including accession numbers GCST0001391 to GCST0002121(Table 1). This dataset comprises information from 563,085 European samples and approximately 22 million SNPs[31].

**Table 1.**
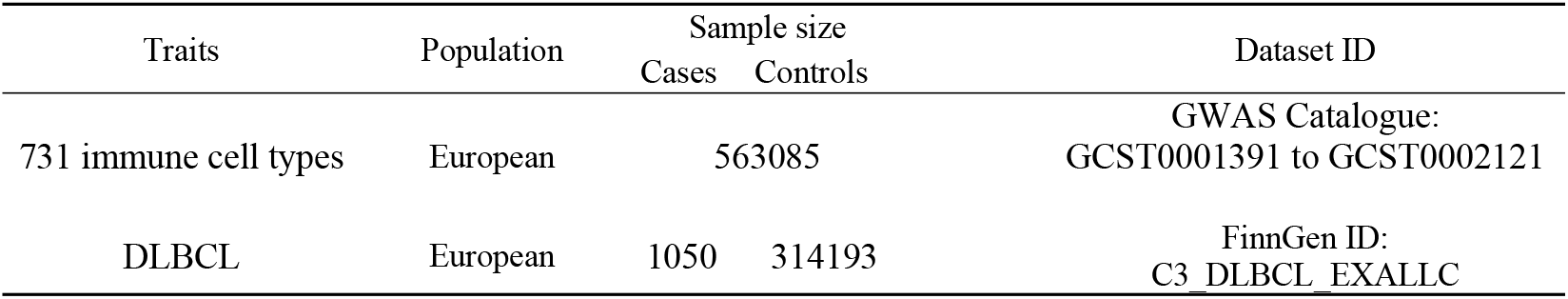
Sources and Information on Relevant Exposures and Outcomes.

Genetic data on DLBCL was sourced from FinnGen R10 (https://r10.finngen.fi/), specifically from the GWAS dataset ID C3_DLBCL_EXALLC (https://storage.googleapis.com/finngen-public-data-r10/summary_stats/finngen_R10_C3_DLBCL_EXALLC.gz). This dataset includes 1,050 DLBCL cases and 314,193 controls from European samples.

### 2.3. Selection of instrumental variables

In this MR analysis, SNPs were used as instrumental variables (IVs) to investigate causal relationships between immune cell characteristics and DLBCL[32]. The selection of IVs adhered to three key assumptions: (1) a strong correlation with the exposure (immune cell traits); (2) no direct association with the outcome (DLBCL); and (3) independence from confounding factors (Figure 1). SNPs were chosen based on a stringent threshold (P < 1 × 10^-5) to ensure relevance[33-35], with parameters r^2=0.001 and kb=10,000 to reduce linkage disequilibrium effects. Only SNPs with an F-statistic greater than 10 were included to avoid bias from weak instruments. Potentially confounding SNPs were identified and excluded using the LDtrait Tool (https://ldlink.nih.gov/?tab=ldtrait)[36].

### 2.4. Statistical analysis

The Inverse Variance Weighted (IVW) method is the standard for MR analysis[37], assuming all instrumental variables are valid and combining Wald ratio estimates to infer causality. In contrast, the weighted median estimator (WME) requires that more than 50% of the IVs are valid SNPs[38]. The WME has the advantage of needing a smaller sample size and offering reduced bias and lower type I error rates compared to other methods. This study utilized both WME and IVW to estimate causal effects, considering results significant if both methods yielded P-values < 0.05, thereby confirming robust causal relationships. Analyses were performed using the TwoSampleMR package in R version 4.3.1.

### 2.5. Sensitivity analysis

Sensitivity analyses were conducted using the Cochran Q test, MR-PRESSO, MR-Egger, and leave-one-out methods. The Cochran Q test and MR-PRESSO assessed heterogeneity among instrumental variables, with P > 0.05 indicating no significant heterogeneity. The MR-Egger regression method was used to test for horizontal pleiotropy, where P < 0.05 for the intercept term suggested its presence. The leave-one-out method was employed to determine if any individual SNP had an undue impact on the association.

### 2.6. Reverse MR Analysis

In our study of DLBCL as an exposure factor, we initially set the SNP association threshold at p = 5e-08, later adjusting to p = 5e-06 due to insufficient SNPs at the former level. Applying screening criteria of p = 5e-06, r2 = 0.001, and kb = 10000, we identified significant SNPs associated with DLBCL. We then investigated the reverse causal relationship between DLBCL and immune cells using methods including IVW, MR-Egger, WME, WM, and Simple Mode.

## 3. Results

### 3.1 Exploration of the causal effect of immunophenotypes on DLBCL

To explore the causal effects of 731 immune cells on DLBCL, we conducted a two-sample MR analysis. Using WME and IVW methods, we identified 17 immune cells significantly associated with DLBCL onset (P < 0.05) (Figure 2).

**Figure 2:**
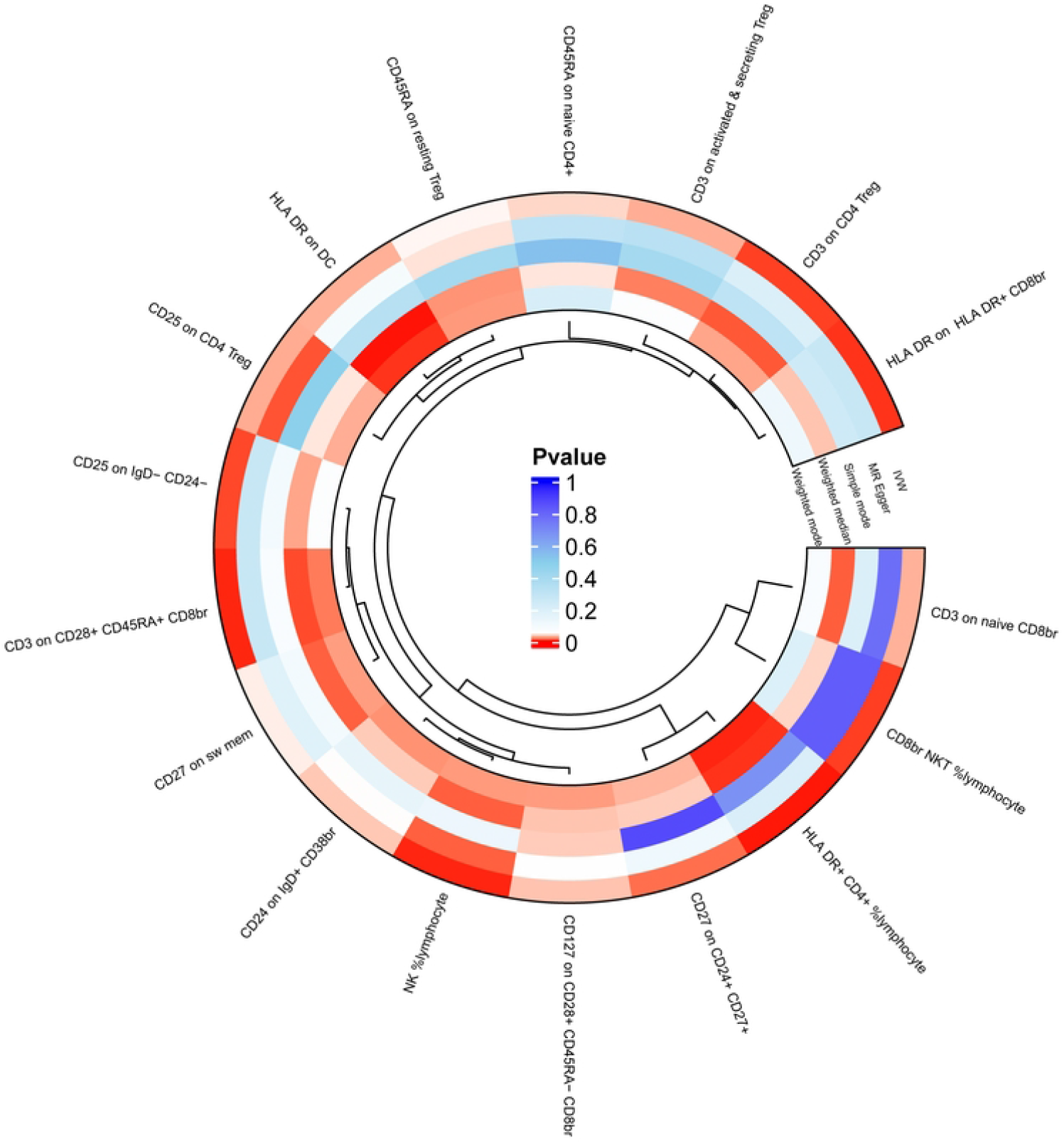
Impact of 17 Immune Cells on DLBCL.

**Figure 3:**
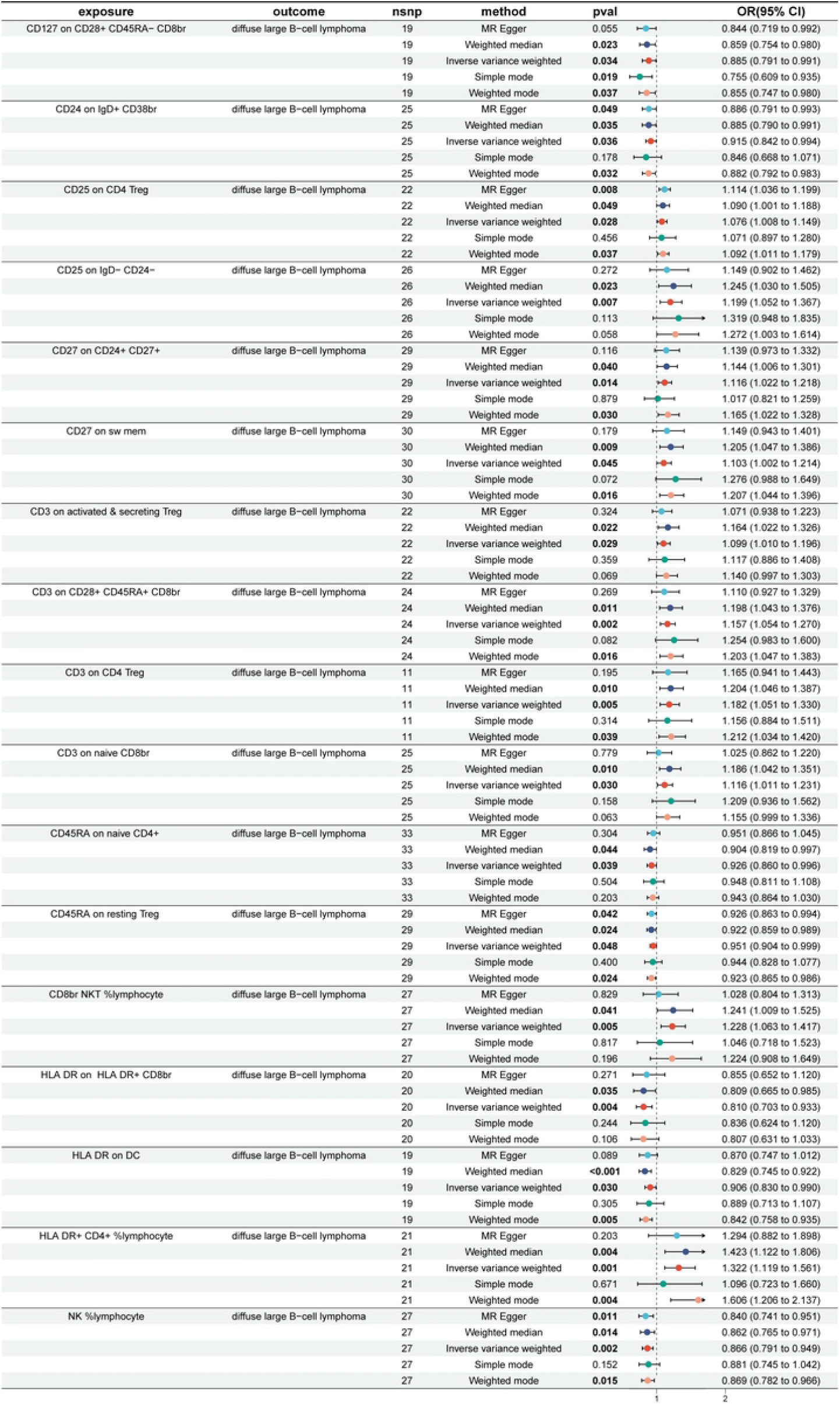
Impact of 17 Immune Cells on DLBCL (WME and IVW : p < 0.05). nsnp, number of single nucleotide polymorphisms; OR, odds ratio; CI, confidence interval.

Seventeen immune features with causal relationships to DLBCL are distributed among different cellular groups: four within the B cell group, two within the maturation stages of the T cell group, six within the Treg group, four within the TBNK group, and one within the dendritic cell (DC) group. The scatter plot illustrating their causal relationship is presented in Supplementary PDF1.

In the B cell group, the protective factor against DLBCL is “CD24 on IgD+ CD38br “(WME: OR: 0.885, p = 0.035; IVW: OR: 0.915, p = 0.036). In contrast, three types of B cells are identified as risk factors for DLBCL:CD25 on IgD-CD24-(WME: OR : 1.245, p =0.023 and IVW: OR : 1.199, p = 0.007), CD27 on CD24+ CD27+ (WME: OR : 1.144, p =0.040 and IVW: OR : 1.116, p = 0.014), and CD27 on sw mem (WME: OR : 1.205, p =0.009 and IVW: OR : 1.103, p = 0.045).

In the maturation stages of T cells, the protective factor is “CD45RA on naive CD4+”(WME: OR : 0.904, p =0.044 and IVW: OR : 0.926, p = 0.039), whereas the risk factor is “CD3 on naive CD8br”(WME: OR : 1.186, p =0.010 and IVW: OR : 1.116, p = 0.030).

In the Treg group, two types are identified as protective factors against DLBCL:CD127 on CD28+ CD45RA-CD8br (WME: OR : 0.859, p =0.023 and IVW: OR : 0.885, p = 0.034), and CD45RA on resting Treg (WME: OR : 0.922, p =0.024 and IVW: OR : 0.951, p = 0.048). Additionally, four types are identified as risk factors for DLBCL:CD25 on CD4 Treg (WME: OR : 1.090, p =0.049 and IVW: OR : 1.076, p = 0.028), CD3 on activated & secreting Treg (WME: OR : 1.164, p =0.022 and IVW: OR : 1.099, p = 0.029), CD3 on CD28+ CD45RA+ CD8br (WME: OR : 1.198, p =0.011 and IVW: OR : 1.157, p = 0.002), and CD3 on CD4 Treg (WME: OR : 1.204, p =0.010 and IVW: OR : 1.182, p = 0.005).

In the TBNK group, two types are identified as protective factors against DLBCL:HLA DR on HLA DR+ CD8br(WME: OR : 0.809, p =0.035 and IVW: OR : 0.810, p = 0.004), and NK %lymphocyte(WME: OR : 0.862, p =0.014 and IVW: OR : 0.866, p = 0.002).

Additionally, two types are identified as risk factors for DLBCL:CD8br NKT %lymphocyte(WME: OR : 1.241, p =0.041 and IVW: OR : 1.228, p = 0.005), and HLA DR+ CD4+ %lymphocyte(WME: OR : 1.423, p =0.004 and IVW: OR : 1.322, p = 0.001).

Lastly, in the DC group, the protective factor against DLBCL is identified as “HLA-DR on DC”(WME: OR : 0.829, p <0.001 and IVW: OR : 0.906, p = 0.030).

### 3.2. Sensitivity analysis

Cochran Q test results revealed no heterogeneity among SNPs (Supplementary Table 1). The MR-Egger intercept test indicated that horizontal pleiotropy did not affect the results (Supplementary Table 2). The funnel plot demonstrated symmetrical distribution, suggesting no bias (Supplementary PDF 2). The MR-PRESSO global test identified no outlier SNPs (Supplementary Table 3). Sensitivity analysis using the leave-one-out method confirmed that individual SNPs did not influence the results (Supplementary PDF 3).

### 3.3. Causal Effects of DLBCL on the four immune cells

The reverse causal analysis using the IVW method revealed a causal relationship between DLBCL and one of the 17 identified immune cells(Figure 4):NK %lymphocyte (IVW: OR: 0.947, p = 0.022).

**Figure 4:**
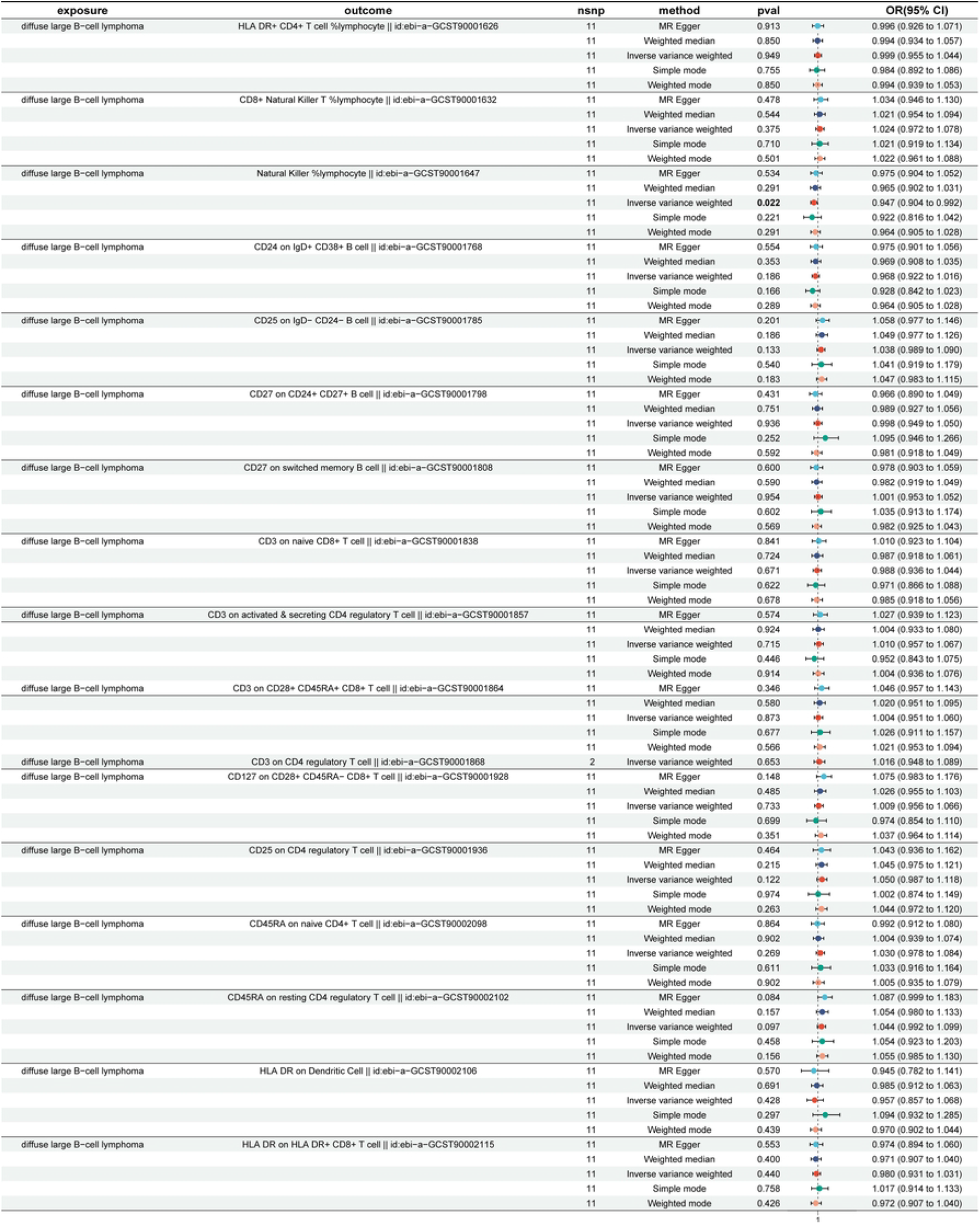
DLBCL and the 17 immune cells. nsnp, number of single nucleotide polymorphisms; OR, odds ratio; CI, confidence interval.

## 4 Discussion

We explored the causal links between 731 immune cell traits and DLBCL using comprehensive public genetic datasets. The WME and IVW methods identified 17 immune cell traits with a significant association to DLBCL. Among them, there are four in the B cell group, two in the maturation stages of the T cell group, six in the Treg group, four in the TBNK group, and one in the dendritic cell group.

In the B cell group, “CD27 on CD24+ CD27+” and “CD27 on sw mem” are considered risk factors for DLBCL. CD27 is a co-stimulatory molecule and a member of the TNF receptor superfamily[39]. CD27, found exclusively on lymphoid cells, is associated with the functional differentiation of T and B cells[40-42]. CD27, upon activation by its ligand CD70, boosts immunoglobulin production in B cells, and sustained CD27 stimulation may contribute to lymphomagenesi[43,44]. Increased serum levels of sCD27 have been reported in acute lymphoblastic leukemia, chronic lymphocytic leukemia, and malignant lymphoma[45,46]. Goto et al.[47] demonstrated that elevated circulating levels of sCD27 in DLBCL patients are associated with a worse prognosis compared to lower levels.

In the Treg group, “CD25 on CD4 Treg” is considered risk factors for DLBCL. CD4+CD25+ regulatory T (Treg) cells, defined by the coexpression of CD4 and CD25, are a subset of suppressor T cells with the ability to inhibit autoreactive T cells[48]. DLBCL patients exhibited an elevated proportion of CD4+CD25+ Treg cells[49]. Circulating sCD25 competes with IL-2 for binding to IL-2R, leading to the suppression of lymphocyte proliferation and reduced NK cell activity, ultimately diminishing overall immune function. Thus, the level of circulating sCD25 serves as a marker for the extent of immune inhibition[50]. Bien and Balcerska[51] propose that the elevated levels of sCD25 in most patients with hematological malignancies are primarily due to tumor cells persistently secreting high amounts of sCD25. Patients with B cell non-Hodgkin lymphoma exhibit significantly elevated levels of sCD25 compared to healthy individuals[52]. Epirubicin, a frequently used chemotherapeutic agent for DLBCL, may enhance immune function by inhibiting sCD25 secretion from Treg cells in affected patients[53].

In the TBNK group, “NK %lymphocyte” is considered a protective factor for DLBCL. NK cells release cytotoxic granules like perforin and granzymes, and produce inflammatory cytokines such as interferon-γ (IFN-γ) and tumor necrosis factor (TNF). While their primary role in antitumor immunity involves direct cytotoxicity, IFN-γ from NK cells is crucial for the efficacy of immunotherapies and for combating lymphoma and other cancers [54-57]. Several cancer therapies, including rituximab, an anti-CD20 monoclonal antibody for B-cell malignancies, and emerging immunotherapeutic approaches like bispecific or trispecific antibodies [58-60], depend on NK cell function. Consequently, the efficacy of various blood cancer treatments hinges on the functional integrity of NK cells. The NK cell population is crucial for immune surveillance and constitutes a key component of innate immunity against virus-infected and malignant cells [61]. NK cells, known for their potent cytotoxicity against tumor cells, significantly contribute to the effectiveness of rituximab-based therapies through their role in antibody-dependent cell cytotoxicity (ADCC) [62,63]. Enhancing NK cell function in these patients may potentially improve the efficacy of lymphoma-targeted antibodies[64]. A low baseline peripheral blood NK cell count correlates with adverse clinical outcomes in patients with DLBCL and follicular lymphoma[65-67]. In DLBCL patients, a reduced NK cell count at diagnosis is associated with decreased survival[67]. This may be linked to cytokine imbalances, as a Phase I study[68] demonstrated that in vivo IL-2 stimulation of NK cells can improve the efficacy of rituximab in treating B-cell lymphomas.

In our study, we performed a two-sample MR analysis using GWAS datasets of approximately 231,771 individuals, ensuring robust statistical power. Genetic instrumental variables were analyzed with a significance threshold of P<0.05 across both WME and IVW methods, enhancing result reliability by mitigating potential issues like horizontal pleiotropy and confounding factors. Older DLBCL patients often have poor baseline health and limited tolerance to immunochemotherapy[69]. Personalized treatment for this group is an unmet need, potentially addressable through targeting new molecular features such as oncogenic mutations and tumor microenvironment. By identifying 17 distinct immune cell types, our research elucidates the intricate interplay between the immune system and DLBCL, offering profound insights into the tumor microenvironment. The comprehensive profiling of immune cell types in our study provides a foundation for novel strategies that target these molecular features, potentially improving outcomes for this vulnerable patient population. Further research should focus on the development and clinical testing of these targeted therapies to validate their efficacy and safety in older or other DLBCL patients.

However, this study has several limitations when interpreted through Mendelian randomization. Firstly, certain immune cell phenotypes remain insufficiently researched, leaving their immunological roles in DLBCL unclear. Secondly, the patient cohort predominantly consists of European individuals, limiting the generalizability of the findings across different geographic populations. To enhance the applicability and understanding of these results, future research should aim to include more diverse populations and further investigate the specific immune cell phenotypes involved in DLBCL.

## 5 Conclusions

Our MR analysis identified seventeen distinct immune cell types, clarifying the complex interactions between the immune system and DLBCL. These findings offer profound insights into the tumor microenvironment and provide a foundation for developing novel immunotherapeutic strategies targeting these molecular features.

## Ethics

This study used only published GWAS datasets, with all original studies having received the necessary ethical approval. Therefore, separate ethical approval was not considered necessary for this review.

## Consent for publication

All authors are aware of and agree to publication without the need for a separate consent form.

## Data Availability

The article and supplementary material contain the original contributions described in the study. If you have any further questions, please contact the corresponding author.

## Conflicts of Interest

The authors declare that they have no competing financial interests.

## Funding Statement

No funding was secured for this study.

## Acknowledgements

We acknowledge the use of summary data on SNPs associated with 731 immune cells and DLBCL from the GWAS Catalogue and FinnGen Biobank, and we extend our gratitude to all participants for their contributions.

## Declarations

Ethics approval and consent to participate

Not applicable. This study was based exclusively on published GWAS datasets, all of which had obtained the necessary ethical approvals and participant consent in the original studies.

Consent for publication Not applicable.

Competing interests

The authors declare no competing interests.

## Notes

### Competing Interest Statement

The authors have declared no competing interest.

